# Idiosyncratic learning performance in flies generalizes across modalities

**DOI:** 10.1101/2021.01.23.427920

**Authors:** Matthew Smith, Kyle S. Honegger, Glenn Turner, Benjamin de Bivort

## Abstract

Individuals vary in their innate behaviors, even when they have the same genome and have been reared in the same environment. The extent of individuality in plastic behaviors, like learning, is less well characterized. Also unknown is the extent to which intragenotypic differences in learning generalize: if an individual performs well in one assay, will it perform well in other assays? We investigated this using the fruit fly *Drosophila melanogaster*, an organism long-used to study the mechanistic basis of learning and memory. We found that isogenic flies, reared in identical lab conditions, and subject to classical conditioning that associated odorants with electric shock, exhibit clear individuality in their learning responses. Flies that performed well when an odor was paired with shock tended to perform well when other odors were paired with shock, or when the original odor was paired with bitter taste. Thus, individuality in learning performance appears to be prominent in isogenic animals reared identically, and individual differences in learning performance generalize across stimulus modalities. Establishing these results in flies opens up the possibility of studying the genetic and neural circuit basis of individual differences in learning in a highly suitable model organism.

## Introduction

Genetically identical Drosophila melanogaster flies display variability in their behaviors even when reared in identical environments [1–8]. Such individuality has been observed in phototaxis [4], spontaneous locomotor biases [3], thermal preference [2], spontaneous microbehaviors [7], and object-fixated locomotion [8]. These differences persist over days and represent something like a fly personality. Work to date has focused exclusively on innate or spontaneous behaviors. But plastic behaviors, such as learning, also have the potential to exhibit individuality, as each animal may have an idiosyncratic propensity to respond to training stimuli [9]. Individual variation in learning within insect populations has been described as early as 1907, by Charles Turner in ants and honey bees [10,11]. To our knowledge, individual variation in learning among genetically identical flies has not been characterized.

Here we present evidence that genetically identical flies exhibit individuality in their ability to learn odor associations. Drawing inspiration from a classical conditioning assay [12–14], animals are exposed to two stimuli, a conditioned stimulus (CS), to which their behavioral response will change across the conditioning, and an unconditioned stimulus (US), to which their response will remain invariant [15]. Typically, the CS is subdivided into two stimuli of the same modality (e.g., two odors) one of which (CS+) is delivered simultaneously with the US, and one of which (CS-) is delivered separately from the US, serving as a control. In a typical *Drosophila* olfactory classical conditioning assay [13], the CS are volatile aversive odorants, often 3-octanol (OCT) and 4-methylcyclohexanol (MCH), and the US is aversive electric shock.

While this assay is normally performed on bulk populations of dozens of flies simultaneously, we individualized it to deliver CS and US to single flies, one per assay chamber in the style of [9]. Our implementation also permitted the selection of different CS odorants under computer control and the use of either electric shock or optogenetic activation [16] of negative valence neural circuit elements as US [17]. This flexibility over the learning stimuli allowed us to ask high-level questions about individuality in learning. For example, does a fly’s performance on one learning task predict its performance on another learning task? We tested this by measuring a fly’s ability to relearn the opposite association to which it was recently trained (i.e., swapping the CS+ and CS-odors). This reversal learning represents a more cognitively demanding form of learning compared to classical conditioning because it requires modification of the previous association. In vertebrates performance on reversal learning tasks has been used as a measure of impulsive and compulsive addiction behaviors [18]. In *Drosophila* studies, reversal learning has been proposed as a metric of cognitive flexibility [19–22]. We also measured the correlation in learning responses between OCT-MCH as CS and an entirely novel odor pair.

Correlated learning performance across multiple CS odors might arise by individual variation in the sensory circuits detecting US cues (an animal that is generally insensitive to shock may fail to learn an odor-shock pairing). This scenario is consistent with the observation that training flies with a higher voltage US leads to better performance in a learning assay as compared to training with a lower voltage US [23]. We examined individual differences in US encoding by training the same flies across multiple US modalities: shock and optogenetic stimulation of bitter taste receptor neurons. We found a positive correlation in the learning response of flies to shock and bitter stimulation. Thus, individual learning performance appears to generalize across odors as well as unconditioned stimuli. This suggests that higher-order plasticity circuits within the brain, rather than the sensory periphery, may harbor the sites where molecular- or circuit-level variation imparts individuality to learning responses. Mapping such “loci of individuality” [24] is a compelling direction for future work to characterize the basis of variation in learning ability in this model system.

## Materials and Methods

All flies were grown on cornmeal/dextrose food in incubators (25C, 40% relative humidity, 12:12 h light:dark cycle). Behavior experiments were conducted females 7-8 days post eclosion. For optogenetic experiments, flies expressing *Gr66a-LexA* in bitter taste receptors were crossed to flies expressing *LexAop-CsChrimson*, the optogenetic activator. Gr66a-LexAp65 was constructed using SLIC cloning [25]. The Gr66a promoter fragment was the same 1798bp segment used in a previous study of Gr66a expression [26], and extended from the translation start site of the Gr66a open reading frame up to the next upstream gene. This was joined to the start codon of the LexA::p65 transcriptional activator from pBPLexA::p65Uw [27] in a vector backbone derived from pUASTattB [28] by removing the UAS sites. The construct was integrated into the attP18 site. In experimental groups receiving the optogenetic US, 10ul of 100mM all-trans-retinal (ATR) was applied to the surface of fly food, and flies were housed on this food for at least 48 hours. Flies were aspirated directly into the behavioral arenas without anesthetization.

The arenas consisted of 15 linear tunnels with inlets at either end and a vent at the center (figure 1A-B). A single fly is placed into each tunnel and is allowed to walk freely. Laminar airflow carrying odor stimuli enters the tunnels from either end and meets at the center, forming a sharp boundary. From there, the odorized air is vented to the room (figure 1B). Odors were delivered by flowing clean air over liquid odorants in a series of vials, under the control of solenoids and mass-flow controllers, as described in [6]. Within the arena, flies were presented one pair of odors (e.g., MCH vs. octanol or 1-pentanol vs. 2-heptanone). During the training phase, both tunnel halves are filled with a single odor, if the odorant is the CS+ then the US is activated when the tunnel is filled with that odorant. For optogenetic experiments, 626nm red LEDs were used to activate csChrimson. These LEDs were pulsed at 20Hz for 3 seconds with a 5 second interstimulus interval. For shock experiments, laser cut sheets of indium-tin-oxide (ITO) were installed in the tunnels to deliver 80V DC pulses from a Grass SD9 Pulse Stimulator at 20Hz for 5 seconds delivered at 10 second intervals. An individual’s learning response was measured by the normalized magnitude of change in occupancy towards the CS-from pre-training to post-training. This metric has a value of 0 if flies exhibit no learning, 1 if they spend all their time post-training in the CS-compartment, and 1 if they spend all their time post-training in the CS+ compartment. Normalizing by the pre-training preference response accounts for individual variation in baseline preference [6].

**Figure 1.**
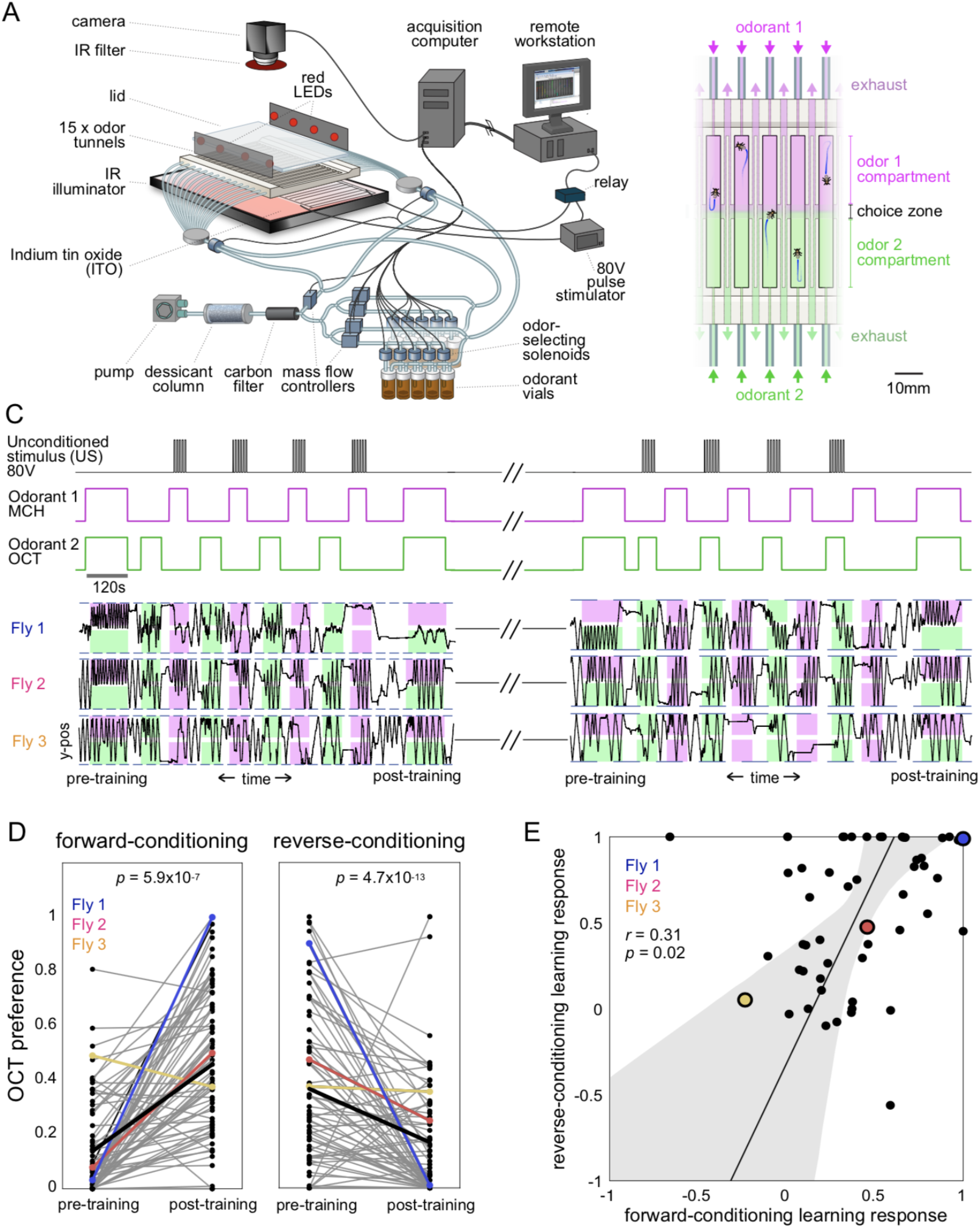
Individuality in associative learning. (A) Schematic of the odor conditioning experimental apparatus. (B) Zoom-in view of the linear behavioral arenas, with odorant flowing into each half. (C) Diagram of training protocol (top). Position in the arena versus time kymographs of three specific flies undergoing conditioning. Magenta and green shading indicate the portions of each arena that are filled with OCT and MCH, respectively. (D) Octanol preference of flies before and after training with MCH as the CS+ (left) and with OCT as the CS+ (right). Points are individual flies. Colored examples correspond to (C). Thick black line represents the mean. (E) Scatterplot of individuals’ learning responses for the reverse-vs forwardconditioning trials (*r* = 0.31; *p* = 0.02; *n* = 53). Points are individual flies. Line is the best linear fit and shaded region is the 95%CI of the best-fit line.

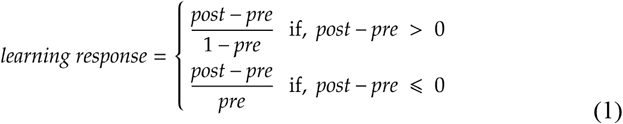

The individual identity of flies whose learning performance was assessed more than once was maintained by housing flies in 96 well plates modified for individual storage (flyPlates, FlySorter, LLC). All learning assays were conducted in a temperature-controlled environmental chamber in total darkness at 25C and 40% relative humidity. Flies were illuminated from below using a modified 15-inch laptop display panel equipped with a high-density infrared LED array for homogenous backlighting, as described in [6]. We used a high-resolution CMOS camera (Point Grey Firefly MV) and longpass filter (Kodak Wratten Filter 87C) to collect 60Hz video. Tracking was performed using custom MATLAB scripts that used 2D cross-correlation for tunnel and initial fly identification, and background subtraction to segment fly boundaries. Data analysis was performed using custom MATLAB scripts.

All raw data and analysis scripts are available at *http://lab.de-bivort.org/individuality-in-learning* and *https://zenodo.org/record/4458572*.

## Results

To characterize the learning responses of individual flies, we built an instrument (figure 1A) that could deliver odor conditioned stimuli and optogenetic and electric shock unconditioned stimuli to 15 flies simultaneously. Laminar airflow carrying odorants originates from the tunnel ends and meets at the center of the tunnel to create a sharp odor-choice boundary (figure 1B), as previously described [6]. We used a classical conditioning and reversal learning protocol (figure 1C) to form associations between odors and the aversive US. In our first experiments we used MCH and OCT (4-methylcyclohexanol and 3-octanol), standard CS odorants for classical conditioning in flies. Each training assay consisted of a 2 min pre-training period in which both odorants were present in the tunnels, allowing us to measure flies′ naïve, untrained odor preference. Flies were then subjected to a series of training blocks, in which the entire tunnel was filled with one odorant or the other, in alternation. Electric shock (US) was triggered when MCH (CS+) filled the entire tunnel. In a final 2 min post-training phase, both odors were presented again. Flies’ preference for OCT was quantified before and after training as the amount of time spent in OCT divided by total time spent in OCT and MCH.

Five minutes after this forward training protocol, flies were subject to the reversal learning protocol, which was structured identically except that OCT was the CS+ odorant. As expected, the classical conditioning and reversal learning assays resulted in significant changes in mean OCT preference across flies (figure 1D). This mean change was not observed in control experiments (e.g., presenting CS only, or breaking its pairing with the US; See figure S1). However, we also observed individual flies that appeared to not learn, with similar preference for OCT pre- and post-training or increased OCT-preference even when OCT was the CS+. These observations could reflect statistical noise, rather than individual variation in learning response. To test this, we examined the correlation between the learning response during the forward-conditioning protocol and the learning response in the reverse-conditioning protocol. This correlation was positive and significant across individual flies (*r* = 0.31, *p* = 0.02), suggesting that individual animals have idiosyncratic learning responses that generalize across the identity of the CS+ odorant. Differences in learning response were not correlated with a fly’s activity (distance traveled) during the assay or initial odor preference (figure S2).

The observation that individual performance following forward- and reverse-conditioning is correlated suggests that learning ability may generalize across modalities in flies. To explore this possibility, we designed a two day assay in which flies were forward- and reverse-conditioned with 1-pentanol and 2-heptanone as CS odors, stored for 24 hours, and forward- and reverse-conditioned with MCH and OCT (figure 2A). In addition, we substituted optogenetic stimulation of bitter taste neurons as the US instead of electric shock. This was done by expressing CsChrimson [16] in bitter taste neurons using a *Gr66a-LexA* driver, and exposing flies to 626nm LED illumination in place of the electric shocks. Replacing shock with bitter taste let us assess whether individuality and correlation between forward- and reverse-conditioning performance is US-specific. In addition, by looking at learning performance after 24 h, we could assess whether individual variation in learning performance is stable over time. As we saw with shock-odor conditioning, flies subject to optogenetic bitter-odor conditioning exhibited mean learned avoidance of the CS+ odor (figure S3) as well as individuality in forward- and reverse-conditioning performance (figure S4). We observed significant correlations in individual learning performance among almost all four conditioning variants in this experiment (0.36 < *r* < 0.59; 10^-5^ < *p* < 3×10^-3^; figure 2B). The exception was MCH+ and 2-heptanone+, for which learning performance was uncorrelated (*r* = 0.06; *p* = 0.61). These results suggest that individuality in learning performance is largely odor CS- and US-independent and stable over at least 24h.

**Figure 2.**
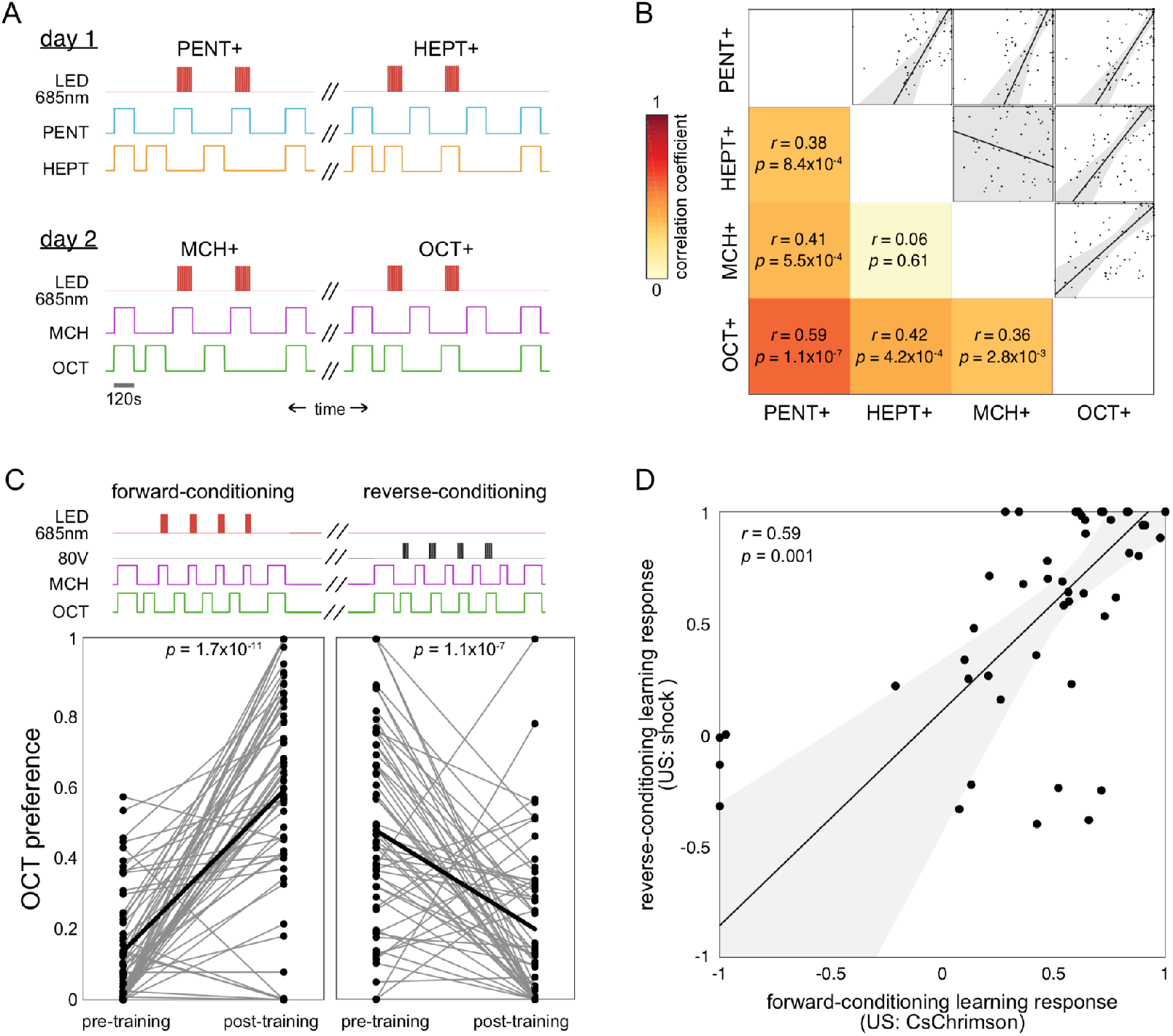
Individual learning across modalities. (A) Schematic of the two day conditioning protocol with two different odor pairs and optogenetic US. (B) Correlation matrix for individual fly learning responses for all pairs of the four different learning trials in (A). x- and y-axes of scatter subplots corre-spond respectively to the learning responses of the CS+ condition indicated by the column and row of the matrix. Points are individual flies. Line is the best linear fit and shaded region is the 95%CI of the best-fit line. (C) Schematic of the back-to-back conditioning paradigms with different US (top). Octanol preference of flies before and after training with shock as the US (left) or optogenetic activation of bitter taste neurons (right). Points are individual flies. Thick black line represents the mean. (D) Scatterplot of learning responses to the shock US trial versus the bitter taste US trial (*r* = 0.45; *p* = 0.01; *n* = 47). Points are individual flies. Line is the best linear fit and shaded region is the 95%CI of the best-fit line.

A possible explanation of these results is individual variation in US encoding. Flies that receive stronger shocks show stronger learning responses [23], so spontaneous variation in the perception of a US (either shock or bitter taste) may affect the learning responses for many CS. We tested this by performing forward- and reverse-conditioning assays with OCT and MCH, but switching between US within the same animals (figure 2C). Both forward- and reverse-conditioning assays show significant mean differences in odor preference (Wilcoxon rank-sum test; *p* = 1.7×10^-9^; figure 2C). Comparing across US modalities, we observed a positive correlation in learning responses (*r* = 0.59; *p* = 1.4×10^-5^; figure 2D). This suggests that in addition to generalizing across CS odorant identity, individual differences in fly learning performance may generalize across US modalities.

## Discussion

Using a training instrument that 1) has versatile control over CS and US and 2) tracks individual learned behavior, we observed that flies are idiosyncratic in their learning performance in classical conditioning paradigms. Flies that perform well for one CS/US pair tend to perform well for other CS and US, suggesting that individual differences in learning performance generalize across stimulus modalities. These results provide a basis upon which to probe the mechanistic basis of individuality in learning. Specifically, our results suggest that the biological basis for such idiosyncrasy in olfactory learning originates more centrally in the brain than sensory circuit elements dedicated to encoding either CS or US. Perhaps there are circuit elements encoding valence generally, which are activated by alternative US and can be paired with alternative CS, that vary stochastically across individuals, accounting for their individual performance [1]. Such sites would be “loci of individuality” [24] for learning performance. Mushroom body dopaminergic neurons (DANs) [29,30] and output neurons MBONs [31] are strong candidates, but valence might also be encoded broadly across multiple populations of higher-order learning circuit neurons [30,32].

Individual bees that performed better in learning assays exhibit greater plasticity in antennal lobe [33] and mushroom body [34] calcium traces. In bumblebees, performance on a visual learning task and the micro glomerular density in the collar of the mushroom body [35] were correlated across individuals. Circuit elements known to exhibit high developmental stochasticity [8] may also be natural locus of individuality candidates. The wiring between projection neurons and mushroom body kenyon cells is highly stochastic [36]. However, that stochasticity may have evolved for the versatile encoding of large odor stimulus spaces, and is distributed over a large population of kenyon cells (~2500). Any single stochastic PN-KC synapse variant may have a vanishing effect amidst such a large pool of (presumably) independent wiring variations. The mushroom body mediates learning and memory for other sensory modalities including vision and taste [37–39]. It is unknown if learning performance generalizes across CS sensory modalities in *Drosophila* (i.e., beyond different kinds of odors). Flies may be a promising model for characterizing the circuit basis of individual variation in generalized learning ability, which is evident even when individuals are genetically identical and reared in the same environment.

## Acknowledgements

We thank Matt Churgin for helpful comments on the manuscript, Ed Soucy, Brett Graham and Joel Greenwood of Harvard’s Center for Brain Science Neuroengineering core for their help with the Grass SD9 Stimulator. B.L.d.B. was supported by a Sloan Research Fellowship, a Klingenstein-Simons Fellowship Award, a Smith Family Odyssey Award, a Harvard/MIT Basic Neuroscience Grant, and National Science Foundation grant no. IOS-1557913. G.C.T. was supported by the Howard Hughes Medical Institute. M.A.-Y.S. was supported by Harvard’s Quantitative Biology Initiative.

## Supplementary figures

**Figure S1.**
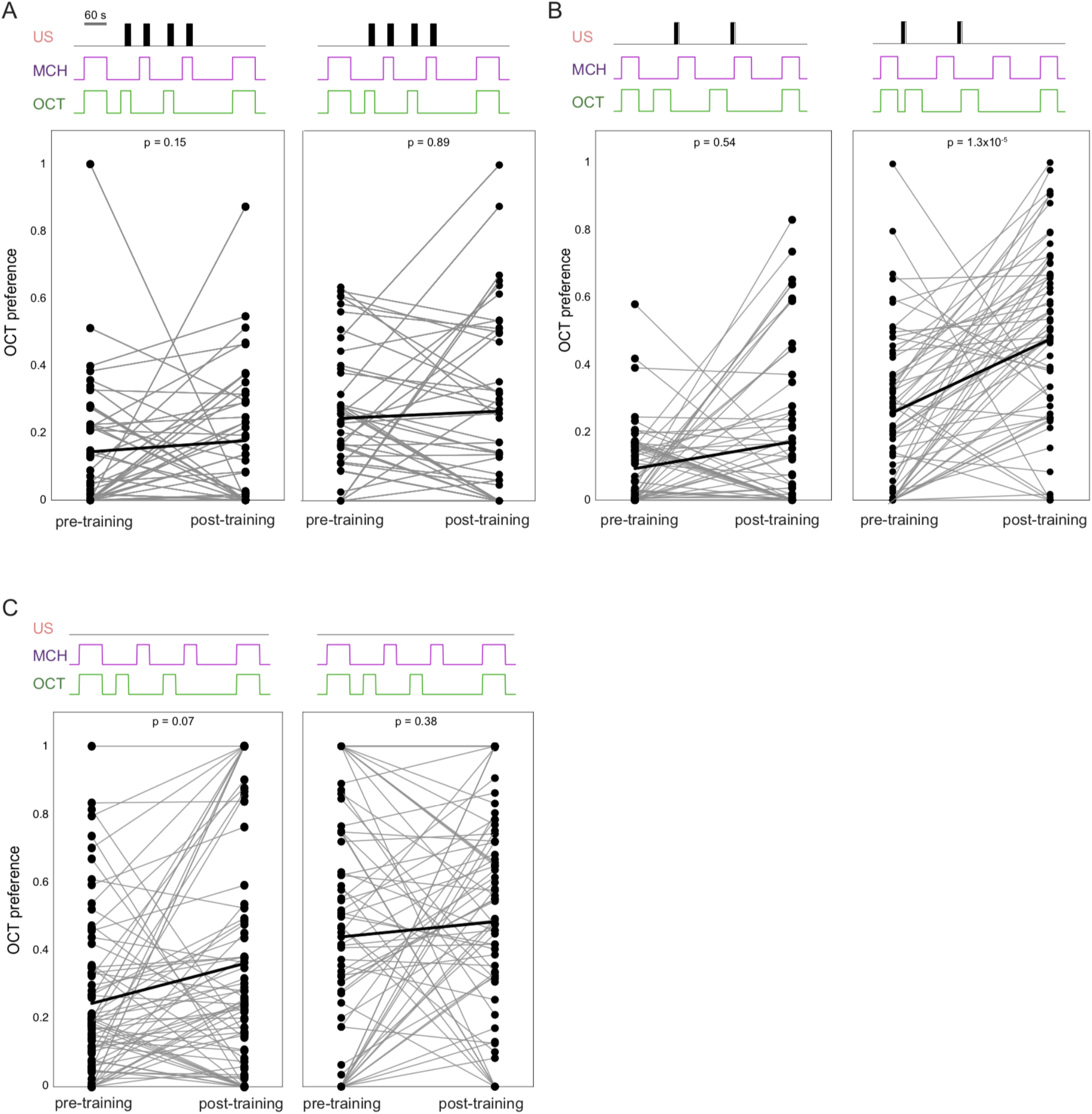
Associative conditioning control experiments. OCT preference of individual flies across pre- and post-contioning periods (left and right dots in each panel, respectively) with two conditioning blocks corresponding to forward- and reverse-conditioning experiments. Dots are individual flies. Thick black line represents mean pre- and post-training. (A) US paired with both OCT and MCH. (B) US presented before the CS. (C) No US presentation. Schematics in style of Figure 1C.

**Figure S2.**
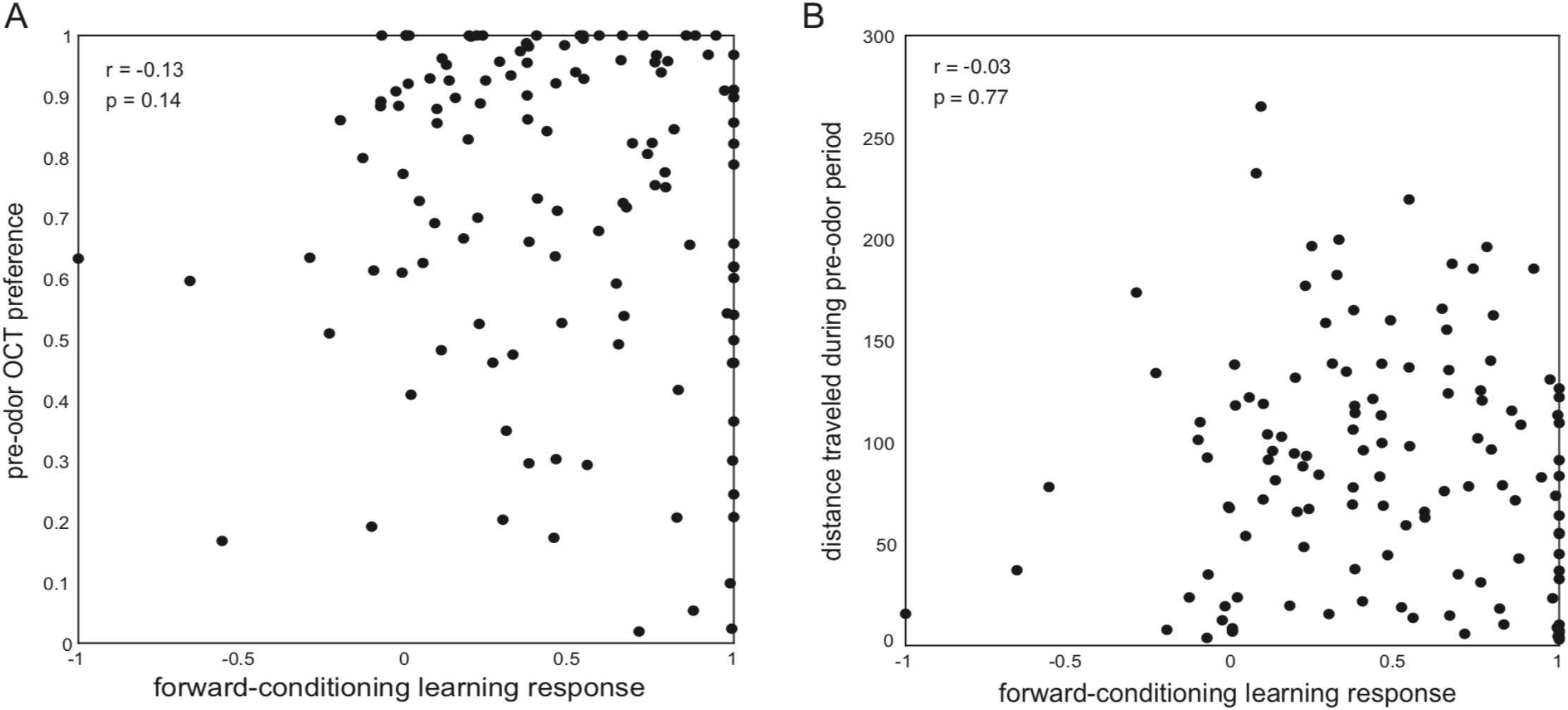
Pre-conditioning behaviors vs learning responses. (A) Scatter plot of individual OCT preference in the pre-conditioning block vs forward-conditioning learning scores (*r* = −0.13; *p* = 0.14). (B) Scatter plot of individual distance traveled during the pre-conditioning block vs forward-conditioning learning score (*r* = −0.03; *p* = 0.77).

**Figure S3.**
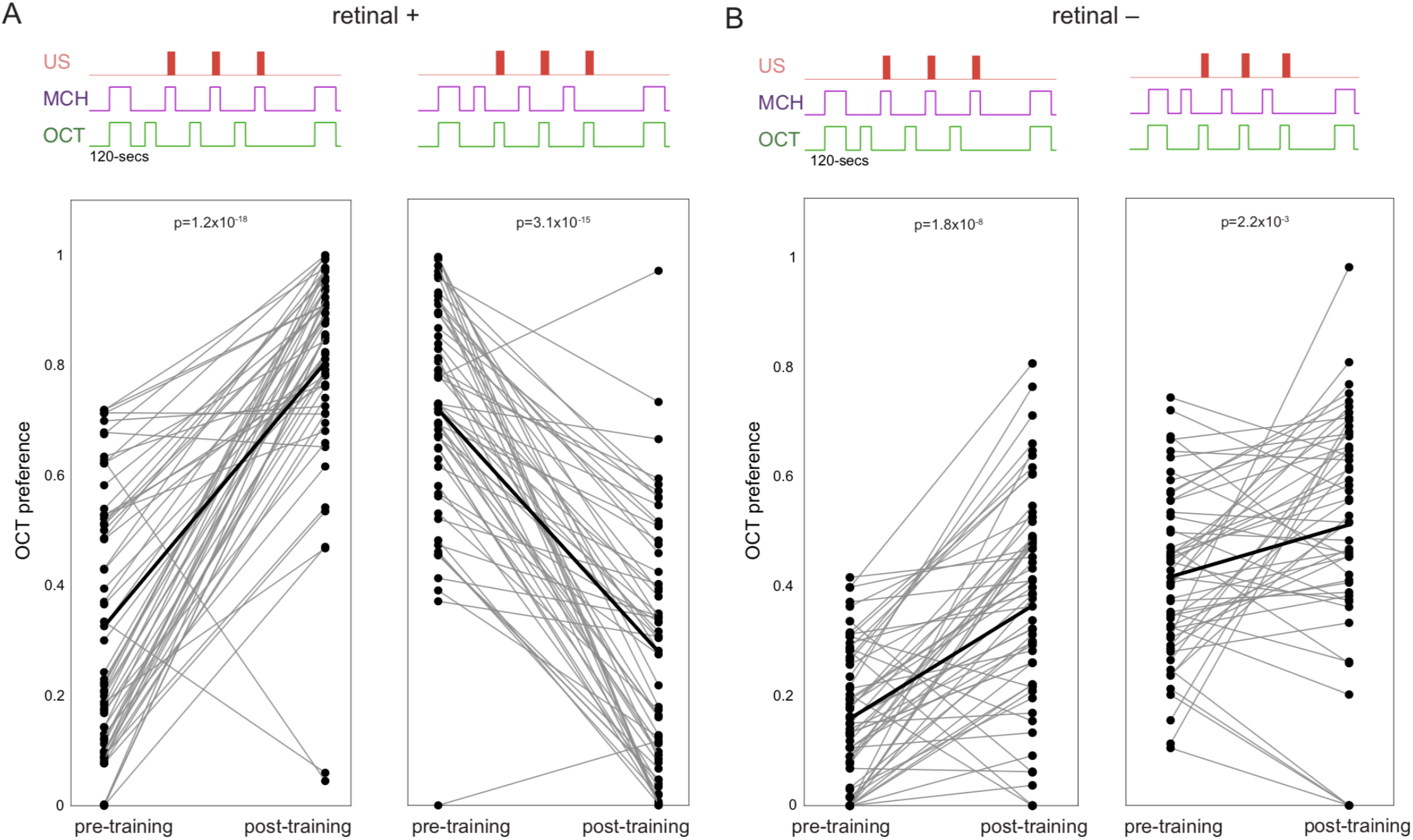
Associative learning with an optogenetic US. (A) OCT preference of *Gr66a-LexA>LexAop-CsChrimson* flies fed all-trans-retinal undergoing forward- and reverse-conditioning. Dots are individual flies. Thick black line represents mean pre- and post-training. (B) OCT preference of *Gr66a-LexA*> *LexAop-CsChrimson* flies not fed all-trans-retinal.

**Figure S4.**
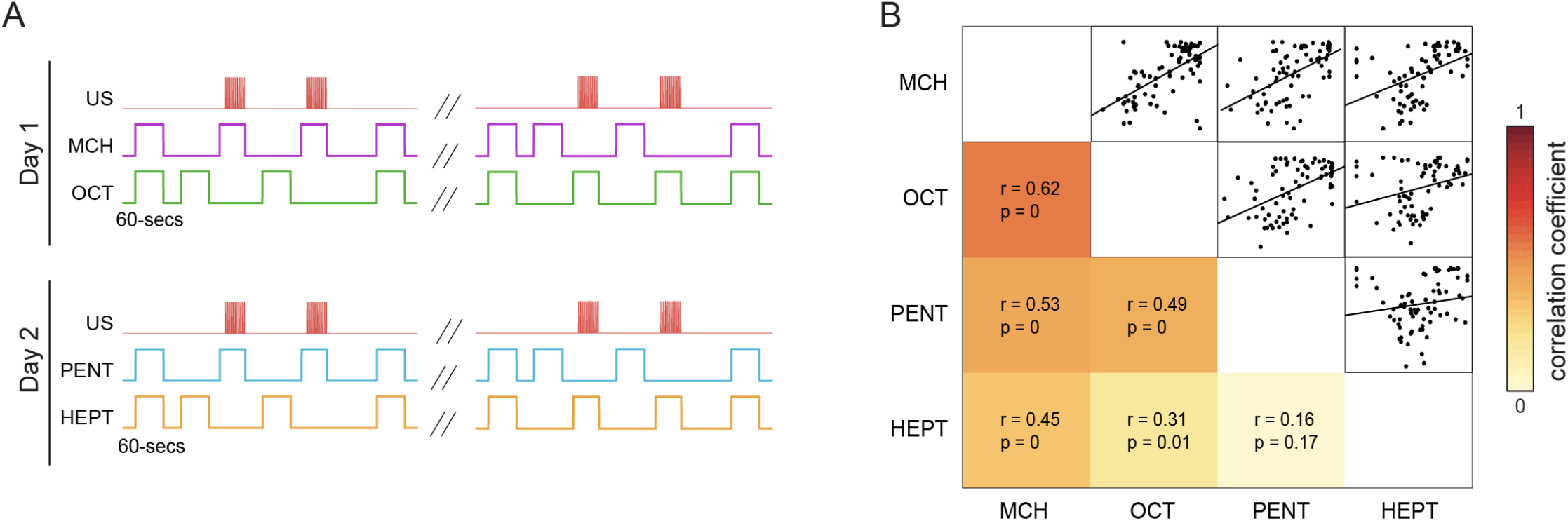
Associative learning across odors. (A) Schematic of the two day conditioning protocol with two different odor pairs and optogenetic US. (B) Correlation matrix for individual fly learning responses for all pairs of the four different conditioning blocks. The MCH-OCT scatterplots shows evidence of individuality in forward- and reverse-conditioning responses. The PENT-HEPT correlation is not significant.

